# Cardiomyocyte-specific adenylyl cyclase type-8 overexpression induces activation of RelA together with myocardial and systemic inflammation

**DOI:** 10.1101/2023.07.15.549173

**Authors:** Vikas Kumar, Kevin Christian Bermea, Dhaneshwar Kumar, Amit Singh, Anjali Verma, Mary Kaileh, Ranjan Sen, Edward G. Lakatta, Luigi Adamo

**Affiliations:** Laboratory of Cardiovascular Science, Intramural Research Program, National Institute on Aging, National Institutes of Health, Baltimore, MD 21224, USA; Laboratory of Molecular Biology & Immunology, Intramural Research Program, National Institute on Aging, National Institutes of Health, Baltimore, MD 21224, USA; Laboratory of Clinical Investigation, Intramural Research Program, National Institute on Aging, National Institutes of Health, Baltimore, MD 21224, USA; Division of Cardiology, Department of Medicine, Johns Hopkins School of Medicine, Baltimore, MD 21205, USA; Immunoregulation Section, Kidney Diseases Branch, National Institute of Diabetes and Digestive and Kidney Diseases (NIDDK), National Institutes of Health, Bethesda, MD, USA

**Keywords:** adenylyl cyclase type 8, accelerated aging, cardiac immunology

## Abstract

**Background:** Mice with cardiac-specific overexpression of adenylyl cyclase (AC) type 8 (TG^AC8^) are under a constant state of severe myocardial stress. They have a remarkable ability to adapt to this stress, but they eventually develop accelerated cardiac aging and experience reduced longevity.

**Results:** Here we demonstrate that activation of ACVIII in cardiomyocytes results in cell-autonomous RelA-mediated NF-κB signaling. This is associated with non-cell-autonomous activation of proinflammatory and age-associated signaling in myocardial endothelial cells and myocardial smooth muscle cells, expansion of myocardial immune cells, increase in serum levels of inflammatory cytokines, and changes in the size or composition of lymphoid organs. These changes precede the appearance of cardiac fibrosis. We provide evidence indicating that ACVIII-driven RelA activation in cardiomyocytes is mediated by calcium-Protein Kinase A (PKA) signaling.

**Conclusions:** Using a model of chronic cardiomyocyte stress and accelerated aging we highlight a novel, PKA/RelA-dependent connection between cardiomyocyte stress, myocardial para-inflammation and systemic inflammation. These findings point to RelA-mediated signaling in cardiomyocytes and inter-organ communication between the heart and lymphoid organs as novel potential therapeutic targets to reduce age-associated myocardial deterioration.

## BACKGROUND

Organisms have developed various adaptations to deal with internal and environmental stress, whether it’s short-term or long-term. As an illustration, when engaging in sudden physical activity, there is an upsurge in autonomic sympathetic-mediated myocardial adenylyl cyclase (AC)/ cyclic AMP (cAMP)/ Protein Kinase A (PKA) /Ca2+ signaling, which plays a pivotal role in responding to both immediate and prolonged stress. In cases of prolonged myocardial stress, such as chronic heart failure and aging, there is a gradual augmentation of autonomic sympathetic signaling and AC/cAMP/PKA/Ca2+ signaling. This escalating signaling contributes to cardiac inflammation and cardiac degeneration. ^1–4^.

There are at least nine different isoforms of AC differing in their biochemical properties and distribution across cells and tissue type^4, 5^. Ca^2+^-calmodulin adenylyl cyclase type 8 (AC8) is not typically expressed in cardiomyocytes. However, constitutively active Ca^2+^-calmodulin adenylyl cyclase type 8 (AC8), driven by the α myosin heavy chain promoter, has been transgenically overexpressed in the murine heart (TG^AC8^) to investigate the relative influence of Ca^2+^ and cAMP on cardiac function, bypassing the potentially deleterious consequences of direct β-Adrenergic receptor stimulation^5, 6^. The TG^AC8^ heart is under constant stress. This is initially associated with “improved performance”^7^, but eventually triggers accelerated aging and premature cardiac dysfunction^6^. This TG^AC8^ transgenic model has therefore been regarded as a useful tool to study both adaptations to stress^7^ and the biology of age-associated cardiac dysfunction^6^.

Bioinformatic analyses of the transcriptome and proteome of left ventricular lysates from young TG^AC8^ mice and wild-type (WT) littermate controls in our previous studies demonstrated that the myocardial-specific overexpression of AC8 is associated with the activation of signaling pathways related to inflammation^7^, before the appearance of signs of premature aging. However, the inflammatory response triggered by cardiomyocyte-specific overexpression of AC8 has not been characterized. Since it is well appreciated that cardiovascular aging is intimately connected with myocardial and systemic immune dysfunction^8^, we performed detailed analyses of the immune response triggered by myocardial-specific activation of constitutive active adenylyl cyclase 8 with the expectation that these analyses would discover mechanistic relationships between chronic cardiomyocyte stress, inflammation, and aging.

## RESULTS

### Cardiac-specific AC8 expression induces myocardial and systemic inflammation

Our group has previously shown that AC8 overexpression is associated with upregulation in the myocardium of several pathways related to inflammation^7^. As an initial step to confirm and expand this observation, we re-analyzed the previously published RNA-seq data (GSEA 4.2.3) from left ventricular (LV) tissue of adult TG^AC8^ mice and age-matched controls. Using FDR value ≤0.05, we identified approximately 2323 genes differentially expressed between TG^AC8^ and WT (Fig. 1A, Supplementary Table 1). To comprehensively investigate the biological meaning of these gene expression changes, we performed PANTHER (protein annotation through evolutionary relationship) pathway analysis. PANTHER pathway analysis confirmed that AC8 activation dysregulated several pathways related to the activation of innate and adaptive immune responses, including “Inflammation mediated by chemokine and cytokine signaling pathway”, “Integrin signaling pathway”, “CCKR signaling map”, “Angiogenesis”, “Gonadotropin-releasing hormone receptor pathway”, “p38-MAPK pathway”, “Angiotensin II-stimulated signaling”, and “Interferon-gamma signaling pathway” (Fig. 1B, Supplementary Table 1). To corroborate this observation, we used Ingenuity Pathway Analysis (IPA) to identify the upstream regulators of the differential gene expression changes observed. IPA analysis confirmed that the majority of the upstream regulators, including *Ccr2, Ifng, Il1b, Ifnar, Tnf,* and *Il2* were directly related to inflammation, cytokine, and chemokine signaling (Fig. 1C, supplementary table 2). Figure 1D shows the increased expression of majority of cytokine, chemokine, and inflammatory signaling genes in the TG^AC8^ heart, compared to the heart of littermate WT animals. As a further validation step, we re-analyzed the proteome profile of TG^AC8^ and WT animals previously reported by our group^7^. This analysis also confirmed the upregulation of inflammatory mediators in the TG^AC8^ mice heart (Supplementary Fig. 1, Supplementary Table 1). This comprehensive analysis of transcriptomic and proteomics data confirmed that overexpression of AC8 in cardiomyocytes is associated with the activation of inflammatory pathways in the myocardium.

**Figure 1.**
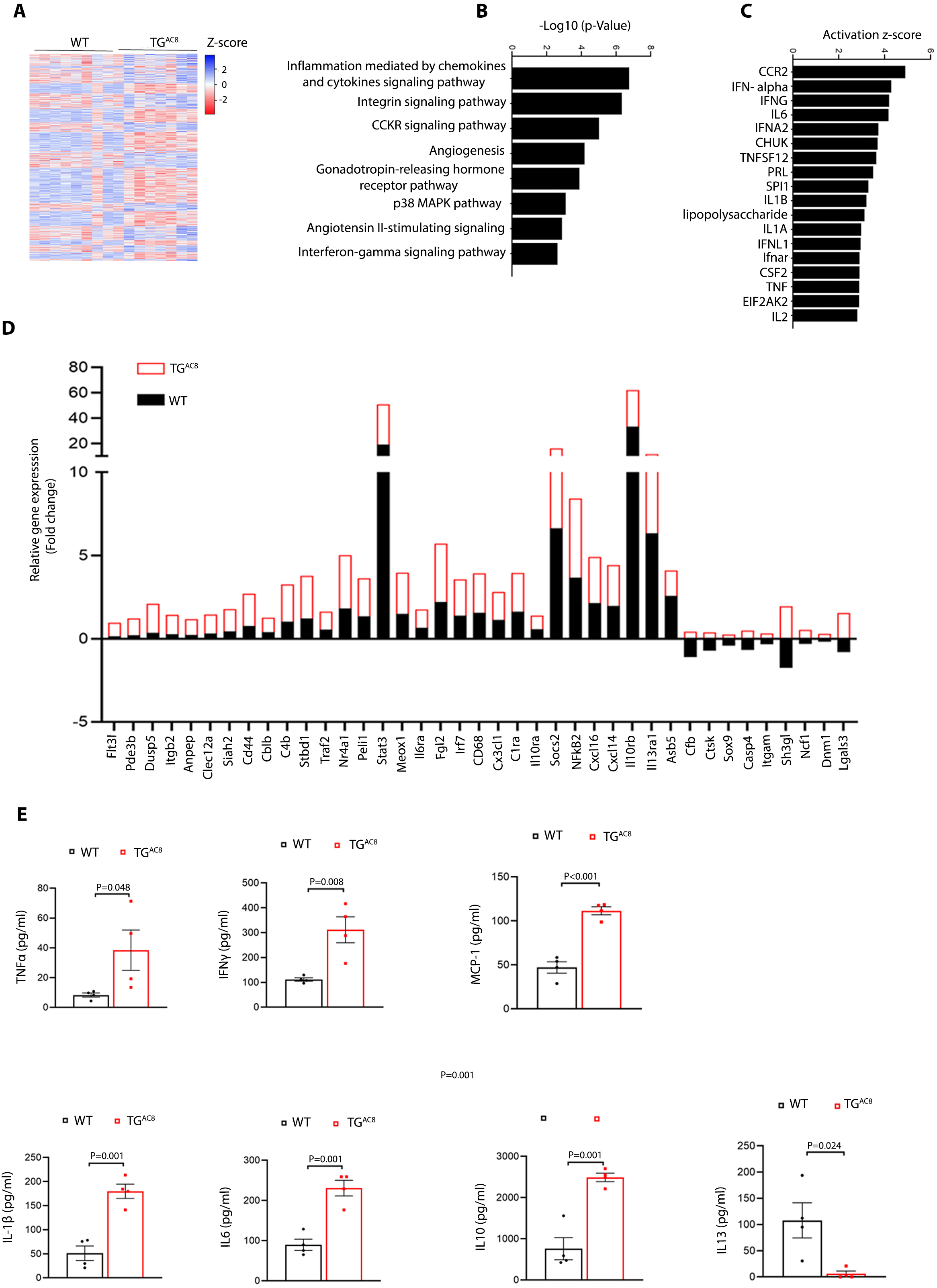
Cardiac-specific AC8 expression induces myocardial and systemic inflammation. **(A)** Heatmap of differentially expressed genes (Fold change>1.5; FDR value<0.05) identified in RNAseq in the left ventricular (LV) tissue between TG^AC8^ and WT mice (n=8 per group). The heatmap was built using the software Heatmapper. **(B)** PANTHER (protein annotation through evolutionary relationship) pathway analysis of differentially expressed genes (Fold change>1.5; FDR value<0.05) in the LV tissue of the heart between TG^AC8^ and WT mice. The top 8 pathways identified are reported. **(C)** Ingenuity Pathway Analysis (IPA) of genes differentially expressed in the LV tissue of TG^AC8^ and WT mice was performed to identify the upstream regulators of the observed gene expression differences. The top 18 regulators, highly enriched in cytokine and chemokine signaling, are reported. **(D)** Relative mRNA expression (RNAseq) of a curated list of cytokine, chemokine, and inflammatory signaling genes differentially expressed between the LV tissue of TG^AC8^ and WT mice (n=8 per group). **(E)** Quantibody inflammation arrays of plasma cytokines and chemokines in TG^AC8^ mice and WT mice (n=8 per group, multiple unpaired t-test).

To understand whether this process was limited to the heart or had systemic effects, we quantified inflammatory cytokines in the plasma using mouse quantibody inflammation arrays. This analysis highlighted higher levels of inflammatory cytokines and chemokines in the plasma of TG^AC8^ animals, including TNFα, IFN-γ, Il1β, IL6, Il10, and MCP-1 (Fig. 1E, Supplementary Table 3). Complete blood count analysis also revealed a higher absolute number of WBCs, lymphocytes, and eosinophils in the TG^AC8^ mice compared to their WT counterparts (Fig. 2A, Supplementary Table 4). Remarkably, the Troponin I assay did not reveal a difference in the serum level of Troponin I in the TG^AC8^ vs WT animals, suggesting that the inflammation was not triggered by cardiac injury (Fig. 2B). The TG^AC8^ heart size was smaller compared to WT mice (Fig. 2C). Trichrome staining analysis did not reveal increased fibrosis in the TG^AC8^ heart, confirming the findings of our prior work^7^ and further suggesting that inflammation was not associated with significant ongoing cardiac damage in these mice (Fig. 2D). Taken together, these results indicate that cardiomyocyte-specific overexpression of AC8 triggers the activation of a myocardial and systemic inflammatory response, that is not driven by cardiac damage, and is not immediately associated with cardiac damage.

**Figure 2.**
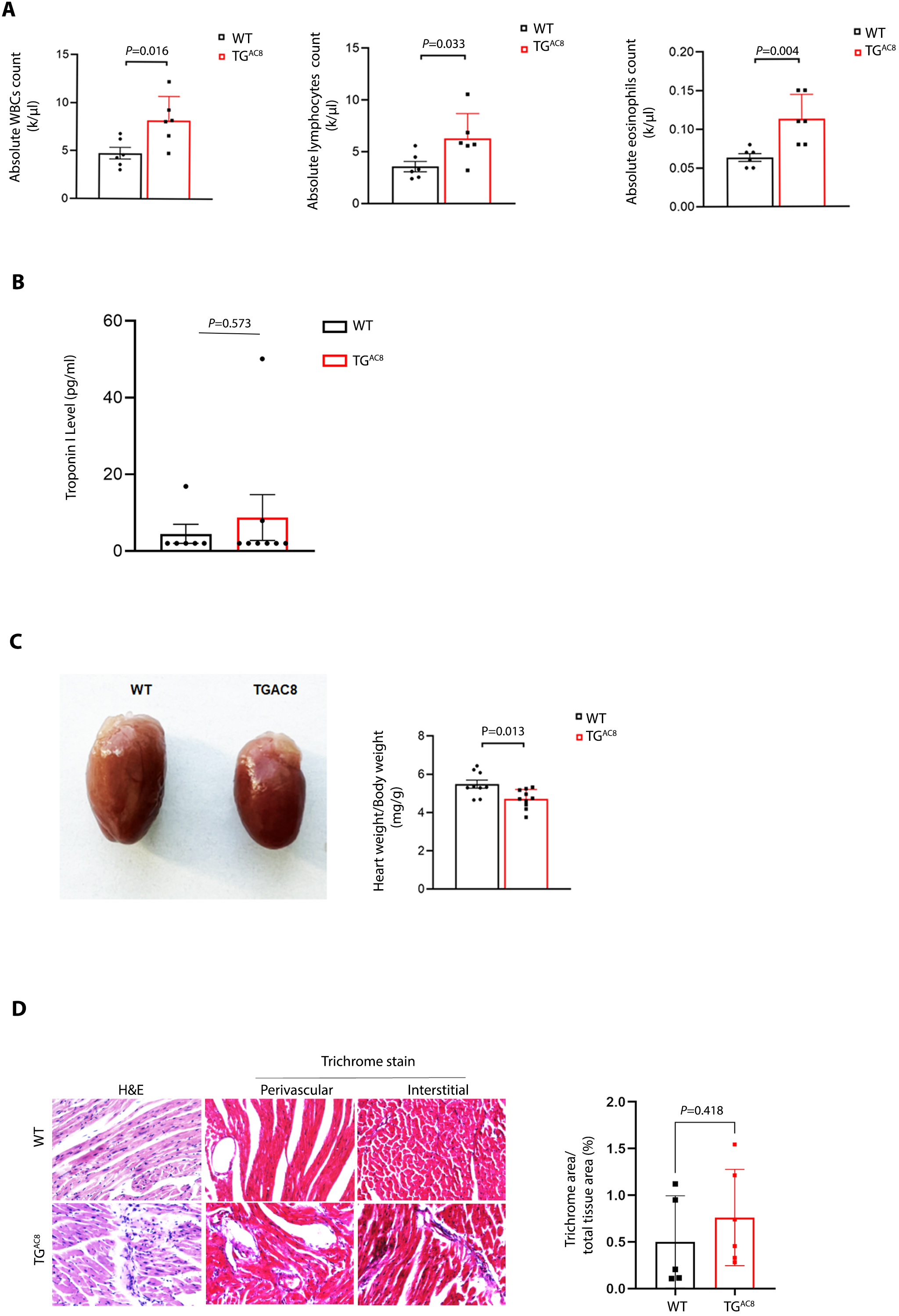
Cardiac-specific AC8 expression-induced inflammation is associated with changes in peripheral blood immune cell counts but is not associated with cardiac injury or fibrosis. **(A)** Automated complete blood count analysis of whole blood from TG^AC8^ mice and WT mice (n=6 per group, multiple unpaired t-test). **(B)** Quantification of Troponin I in plasma of TG^AC8^ mice vs WT mice (n=8-10 per group, multiple unpaired t-test). **(C)** Representative image of the whole heart and quantification of heart weight/body weight in TG^AC8^ mice vs WT mice (n=8-10 per group, multiple unpaired t-test). **(D)** Representative photomicrograph of Hematoxylin & Eosin and Trichrome stained heart tissue section (5um) and quantification of the trichrome positive area in TG^AC8^ mice vs WT mice (n=6 per group, multiple unpaired t-test).

### Cardiac-specific AC8 expression results in the recruitment of multilineage immune cells to the heart

Activation of inflammatory pathways in the heart is typically associated with the expansion and recruitment of immune cells^11, 12^. We, therefore, decided to characterize immune cell populations in the TG^AC8^ heart and WT heart via multicolor flow cytometry and immunohistochemistry. The gating strategy used for the analysis of immune cells by flow cytometry is shown in Supplementary Figure 2. Multi-color flow cytometry revealed that the TG^AC8^ heart was characterized by expansion of myocardial immune cells (CD45+ cells, 44.97% increase, p=0.005). Focusing on specific sub-populations of myeloid cells, we observed an increase in CD45^+^CD11b^+^Ly6G^-^ cells that appeared to be driven mostly by an increase in macrophages (CD64^+^Ly6C^low^ cells, 38.87% increase, p=0.007, Figure 3A-3C). When focusing on cardiac macrophage subpopulations^13, 14^, AC8 overexpression was associated with the expansion of most macrophage subgroups identified by CCR2 and MHCII expression levels. The greatest difference was observed in resident CCR2^-^MHCII^+^ macrophages which increased by 83.53% (p<0.001) (Figure 3D). However, also bone marrow-derived CCR2^+^MHC-II^+^ macrophages increased by 61.33% (p<0.001), and resident CCR2^-^ MHCII^-^ macrophages increased by 59.27% (p=0.002). The number of CCR2^+^ MHCII^-^ macrophages per mg of tissue did not statistically increase in the TG^AC8^ hearts (p=0.172, Figure 3D). We confirmed the increase in macrophages observed in AC8 overexpressing hearts via immunohistochemistry staining (IHC) for CD68 (253.27% increase, p=0.048, Figure 3E). We also analyzed via flow cytometry and IHC cardiac expression of CD19^+^ B cells as well as CD4^+^ and CD8^+^ T cells (Figure 3F-3L). B cells increased by 53.83% (CD19^+^ cells, p=0.051, Figure 3F, 3G) as assessed by flow cytometry and 106.70% (B220^+^ cells, p=0.003) as assessed by IHC (Figure 3H). We also quantified via flow cytometry subclasses of B cells. We found that the AC8 overexpressing heart was characterized by an expansion of CD19^+^ CD11b^+^ B1 cells (61.67% increase, p<0.001). When grouping CD19^+^ CD11b^-^ B2 cells by levels of IgM and IgD expression, we found that overexpression of AC8 was associated with an increase in CD19^+^ IgM^-^IgD^-^ B cells (51.24% increase, p=0.035, Figure 3G) but not in IgM^+^IgD^+^ or IgD^-^IgM^+^ cells. The TG^AC8^ heart was also characterized by expansion of CD4^+^ T cells (51.80%, p=0.018 as assessed by flow cytometry and 79.01%, p=0.049 as assessed by IHC, Figure 3I, 3J) and CD8^+^ T cells (48.47% increase p=0.015 via flow cytometry and 66.06% increase p=0.043 via IHC, Figure 3K, 3L).

**Figure 3.**
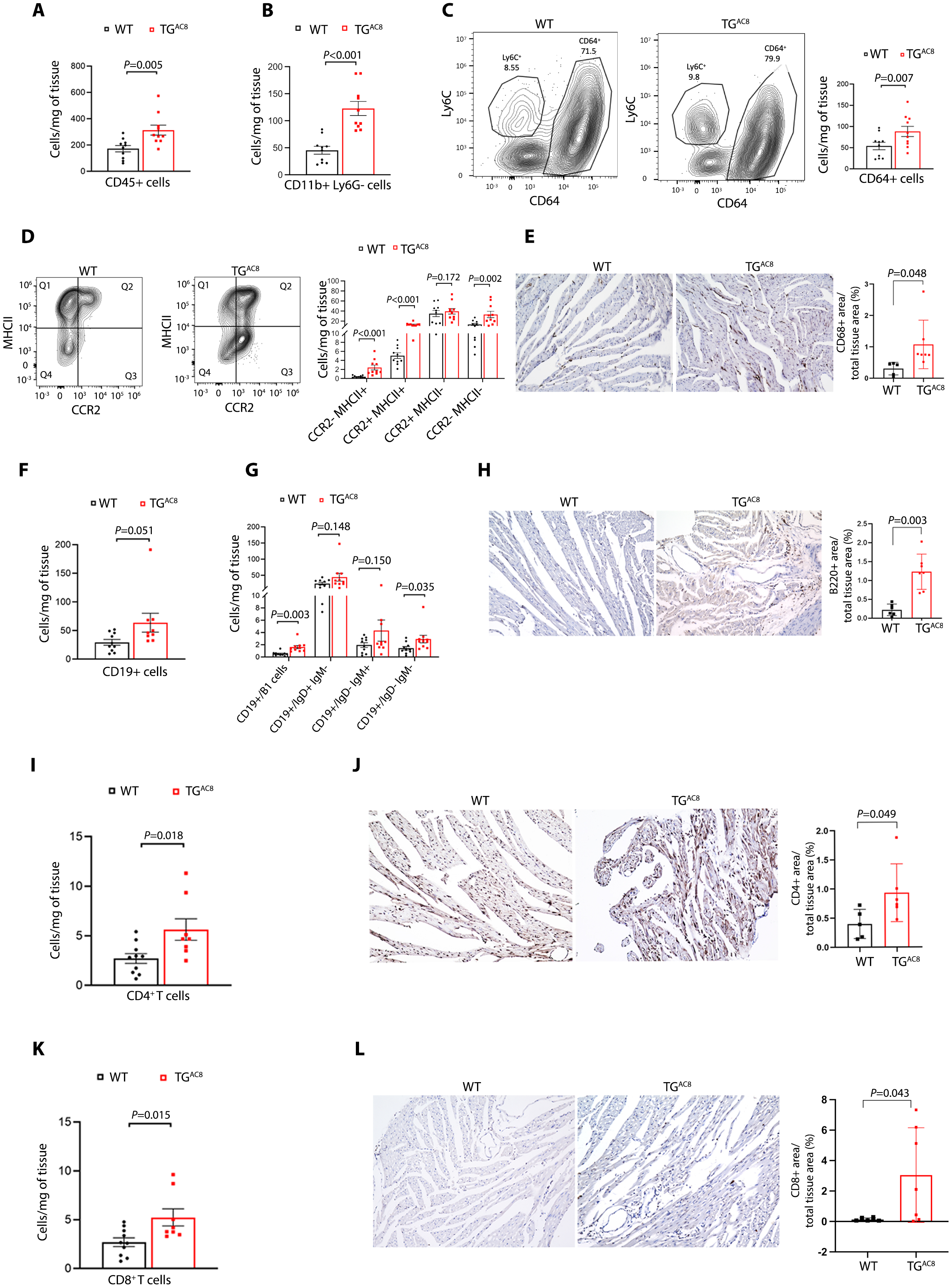
Cardiac-specific AC8 expression results in the recruitment of multilineage immune cells to the heart. **(A), (B)** CD45+ cells and CD45+ CD11b+ Ly6G-cells were identified using multi-color flow cytometry in TG^AC8^ heart vs WT heart (n= 10 per group, multiple unpaired t-test). **(C)** Representative FACS plots and quantification of CD64+ Ly6C^low^ macrophages identified using multi-color flow cytometry in TG^AC8^ heart vs WT heart (n=10 per group, multiple unpaired t-test). **(D)** Representative FACS plots and quantification of macrophage subgroups (CCR2-MHCII+, CCR2+ MHC-II-, CCR2+ MHC-II+, CCR2-MHC-II-macrophages) identified using multi-color flow cytometry in the TG^AC8^ heart vs WT heart (n=8-10, multiple unpaired t-test). **(E)** Representative photomicrograph and quantification of CD68+ macrophages identified using immunohistochemistry (IHC) in TG^AC8^ heart vs WT heart (5uM paraffin sections, n=6 per group, multiple unpaired t-test). **(F), (G)** CD45+ CD19+ B cells and CD19+ B cell subgroups (CD19+ CD11b+ B1cells and, among CD19+ CD11b-B2 cells: IgM+ IgD+, IgM-IgD-, IgM+IgD- and IgM-IgD+ are reported) were identified using multi-color flow cytometry in TG^AC8^ heart vs WT heart (n= 10 per group, multiple unpaired t-test). **(H)** Representative photomicrograph and quantification of B220+ B cells identified using immunohistochemistry (IHC) in TG^AC8^ heart vs WT heart (5uM paraffin sections, n=6 per group, multiple unpaired t-test). **(I)** CD4+ T cell were identified using multi-color flow cytometry in TG^AC8^ heart vs WT heart (n= 10 per group, multiple unpaired t-test). **(J)** Representative photomicrograph and quantification of CD4+ T cells identified using immunohistochemistry (IHC) in TG^AC8^ heart vs WT heart (5uM paraffin sections, n=6 per group, multiple unpaired t-test). **(K)** CD8+ T cell were identified using multi-color flow cytometry in TG^AC8^ heart vs WT heart (n= 10 per group, multiple unpaired t-test). **(L)** Representative photomicrograph and quantification of CD8+ T cells identified using immunohistochemistry (IHC) analysis in TG^AC8^ heart vs WT heart (5µM paraffin sections, n=6 per group, multiple unpaired t-test). Statistics: data is represented as mean± standard error of the mean.

To further characterize the inflammatory response associated with myocardial AC8 overexpression, we analyzed spleen size (Figure 4A) as well as the number and distribution of immune cell populations in the blood, bone marrow, and inguinal lymph nodes (Figure 4 B-J). Myocardial overexpression of AC8 was associated with a reduction in the relative size of the spleen (spleen weight/body weight 25% reduced, P<0.001, Figure 4A). It was also associated with a higher number of CD45+ cells in the blood (59.50% increase, p<0.001) and bone marrow (15.88% increase, p=0.050, Figure 4B), a higher number of monocytes (CD11b+Ly6G-cells) in the blood (33.84% increase, p=0.030) and bone marrow (32.63% increase, p=0.041, Figure 4C, 4D), and a higher number of CD4+ T cells in the blood (43.97% increase, p=0.0015, Figure 4E, 4F). An increase in the number of CD8+ T cells (36.96% increase, p=0.041, Figure 4F, 4G, supplementary figure 2) and CD19+ B cells (64.39% increase, p=0.010; Figure 4H, 4I, supplementary figure 2) was also observed in the peripheral blood of TG^AC8^ mice.

**Figure 4.**
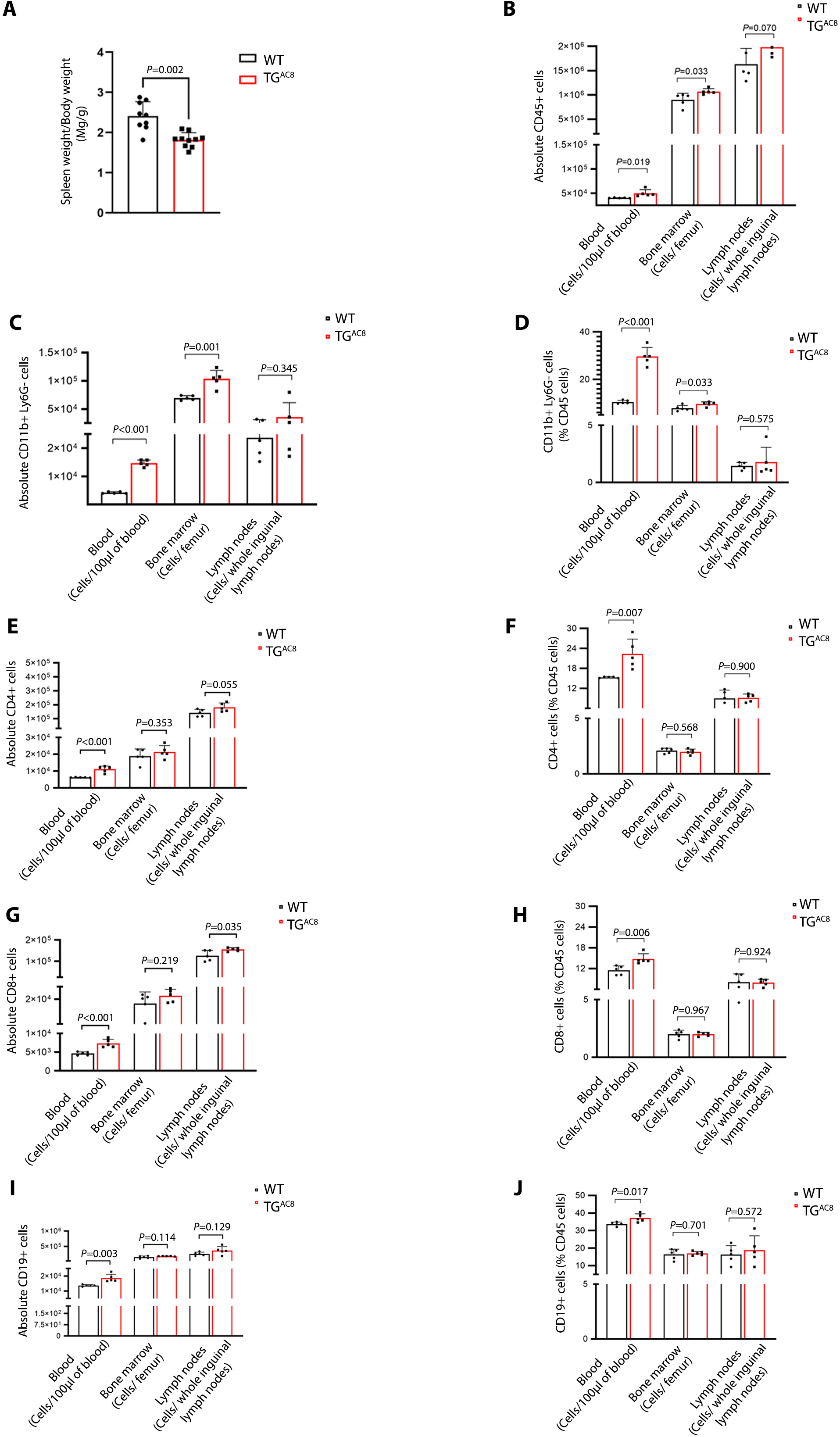
Myocardial AC8 overexpression is associated with a systemic inflammatory response. **A,** Gravimetric analysis of spleens from TG^AC8^ heart vs WT heart, the two groups are compared with unpaired t-test. **B-J**, Flow cytometry-based quantification of immune cells in the blood, bone marrow, and inguinal lymph nodes. The total no. of cells per 100 ul of blood, per femur (1/30 of total cells isolated), or per whole inguinal lymph nodes was quantified via flow cytometry. Data are shown either as an absolute number of cells analyzed (**B, C, E, G, I**) or as % of CD45+ cells within each compartment (**D, F, H, J**). (n= 6 per group, multiple unpaired t-test). Graph bars represent the mean ± Standard Error of the Mean.

### Single-cell RNAseq shows that cardiomyocyte-specific AC8 expression results in the activation of immune signaling pathways in myocardial endothelial cells and smooth muscle cells

To gain further insight into the effects of cardiomyocyte-specific AC8 overexpression on the non-cardiomyocyte component of the myocardium, we used high-throughput single-cell RNA sequencing. We digested hearts from TG^AC8^ animals and WT controls, FACS sorted and pooled equal numbers of endothelial cells (CD31^+^), leukocytes (CD45^+^), and non-endothelial and non-leukocytes (CD31^-^CD45^-^ cells, Figure 5A), and performed high-throughput single-cell RNA sequencing. We first visualized the data using Uniform Manifold Approximation and Projection for Dimension Reduction (UMAP) plots and identified cell types using a recently described automated algorithm that categorizes cells based on specific biomarker combinations^15^ (Figure 5B). We then performed differential gene expression analysis comparing each cell type in AC8 and WT animals and used Ingenuity Pathways Analysis to identify pathways with differential activation within the two experimental conditions. Among nonimmune cells, endothelial cells showed the greatest biological response to cardiomyocyte-specific AC8 overexpression. As shown in Figure 5C, endothelial cells from TG^AC8^ hearts showed upregulation of multiple pathways related to fibrosis as well as activation of endothelin signaling and upregulation of pathways related to cardiac hypertrophy. Smooth muscle cells also showed evidence of activation as shown by marked upregulation of oxidative phosphorylation and evidence of activation of multiple inflammation-related pathways (Supplementary Figure 3). Fibroblasts showed the least number of dysregulated pathways (Supplementary Figure 4). We then looked at the different immune cell types. As shown in Supplementary Figure 5, among immune cells, B cells showed the greatest degree of activation as indicated by broad-ranging activation of several pathways related to cytokine signaling and immune response, but all immune cell types isolated from the TG^AC8^ heart showed evidence of activation when compared to their counterpart isolated from the WT heart (Supplementary Figures 6-11). Across most cell types analyzed, AC8-derived cells showed upregulation of signaling pathways related to immune system activation including leukocyte extravasation signaling and macrophage alternative activation signaling in macrophages (Supplementary Figure 9), CXCR4 and IL3 signaling in glial-like cells (Supplementary Figure 8), HIPPO signaling pathway in CD4 T cells (Supplementary Figure 7), TNFR1 signaling in CD8T cells (Supplementary Figure 6), LPS stimulated MAPK signaling in B cells (Supplementary Figure 5), IL8 signaling in neutrophils (Supplementary Figure 3).

**Figure 5.**
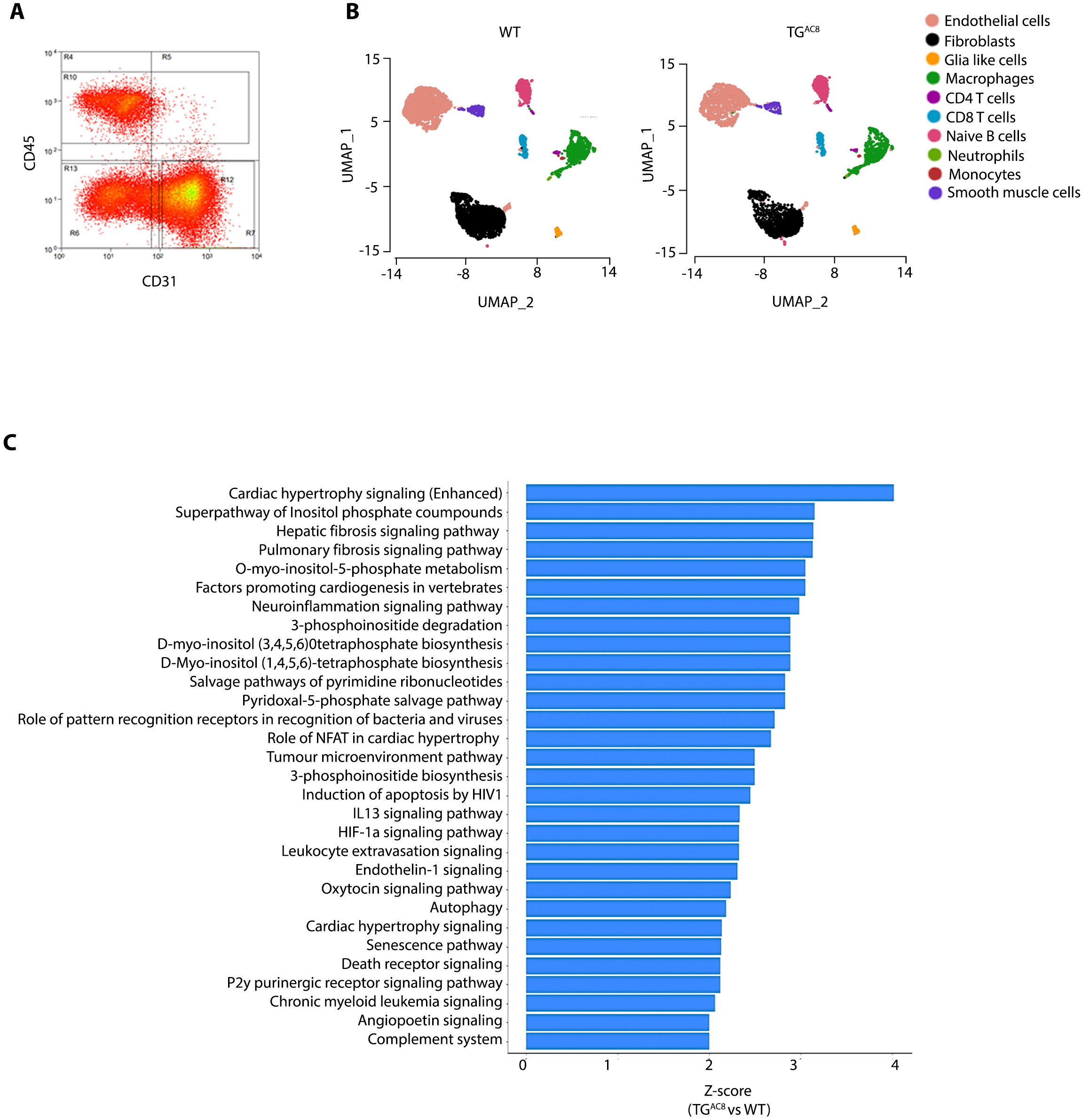
Single-Cell RNAseq shows that cardiomyocyte-specific AC8 expression results in the activation of immune signaling pathways in endothelial cells. **(A)** Non-cardiomyocytes CD31+ CD45-, CD31-CD45+, and CD31-CD45-cells were FACS sorted from TG^AC8^ heart and WT controls and mixed in equal numbers for downstream single-cell sequencing. **(B)** UMAP plot of the single-cell sequencing data of cells sorted from the hearts of WT (left panel) and TGAC8 (right panel) animals. The data was Log2 normalized with a scale factor of 10,000. Outlier identification was made using the variance stabilizing transformation method with 2000 features selected as top variable features. The number of principal components was set at 50 for PCA, followed by a K nearest neighbors search with a k set on 20 and the dimension of reduction to 1:10. Clusters were identified with a resolution of 0.8, and the UMAP plot was created with a dimension of reduction set to 1:10. Finally, cell type was classified using the automated cell-type identification tool described in the main text. **(C)** Ingenuity pathway analysis (IPA) of genes differentially expressed in endothelial cells from TG^AC8^ and WT hearts. Genes with a 1.5 fold-change and p-value < 0.05 were used for pathway analysis. The graph shows the pathways with a z-score > 2 and a p-value < 0.05 in the IPA analysis.

### AC8 overexpression triggers RelA nuclear translocation and activation of inflammatory signaling in cardiomyocytes

The results presented thus far indicate that AC8 overexpression is associated with a remarkable activation of a broad range of inflammatory responses within a variety of myocardial cell types, as well as in the serum, circulating blood, and primary and secondary lymphoid organs. Since our prior work highlighted that AC8 overexpression is associated with marked activation of the AMPK/PKA signaling pathway^7^, we performed a supervised IPA-mediated analysis of genes differentially expressed between TG^AC8^ and WT left ventricular RNAseq datasets, with the specific goal of highlighting potential connections between cAMP, PKA and key regulators of inflammation. This analysis highlighted potential connections between cAMP signaling, PKA signaling, and RelA, a component of the NF-κB signaling complex (Figure 6A). We, therefore, decided to comprehensively assess the activation of NF-κB in the context of AC8 overexpression. As a first step, we assessed the expression in the LV tissue of TG^AC8^ heart of the mRNAs of 5 NF-κB subunits: *RelA, RelB, c-Rel, NF-kB1,* and *NF-kB2*. Among these, *RelA* and *RelB* genes were significantly upregulated in the TG^AC8^ mice heart (Fig. 6B). To confirm these findings, we performed Western blots to assess protein expression levels of RelA, RelB, and c-Rel. Among these, only RelA level increased in the TG^AC8^ heart as compared to WT littermate controls (Fig 6C). Western blot analysis revealed a higher level of RelA in the nuclear lysate of LV tissue of the TG^AC8^ heart and thus confirmed RelA migration into the nucleus (Figure 6D). Further, nuclear lysates of primary cardiomyocytes isolated from TG^AC8^ mice also demonstrated higher nuclear expression of RelA in these cells compared to cells of their WT counterparts (Fig. 6E). These results suggest that AC8 activation in the TG^AC8^ heart is associated with overexpression of RelA and activation of RelA signaling.

**Figure 6.**
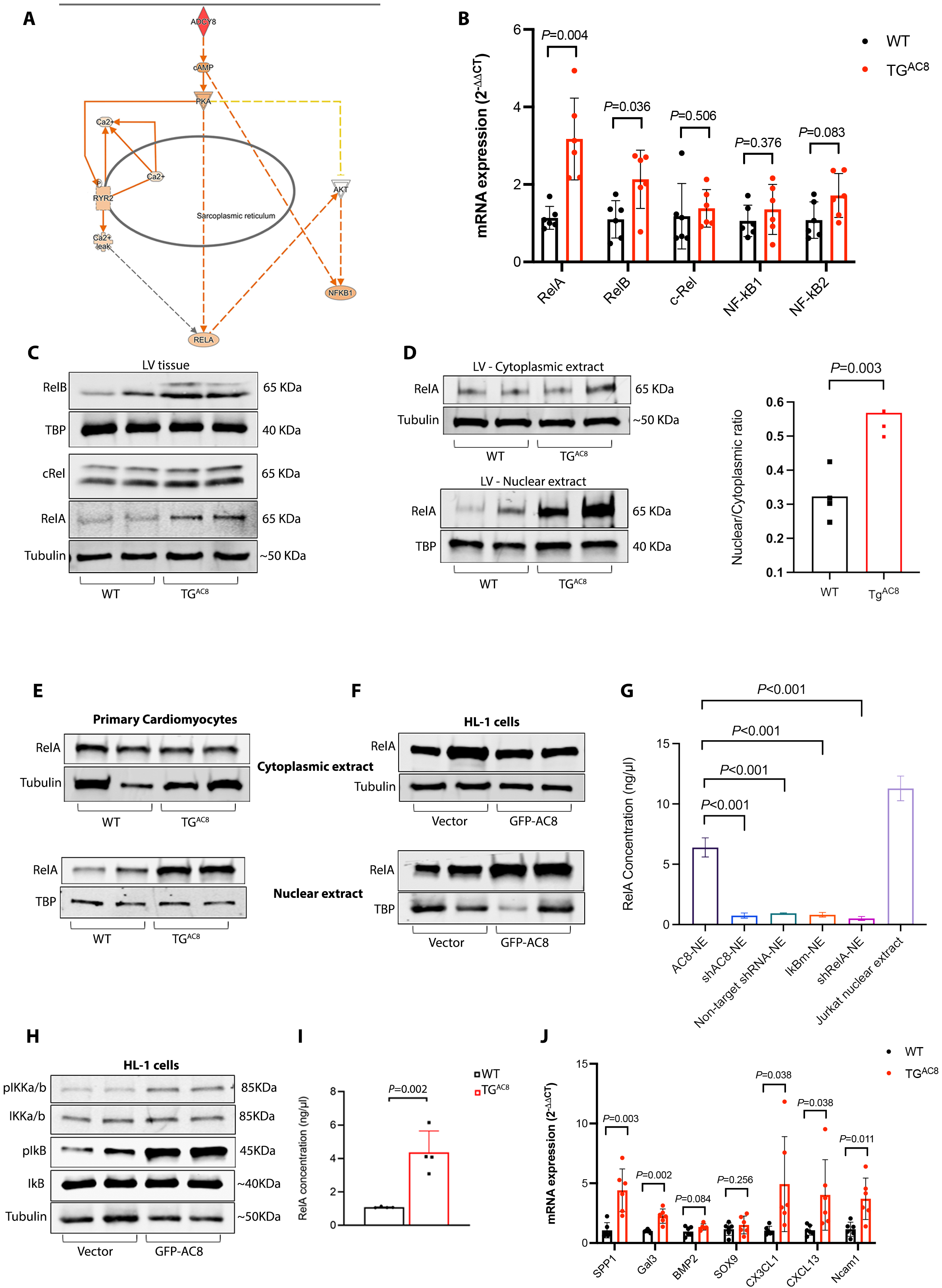
AC8 overexpression triggers RelA nuclear translocation and activation of inflammatory signaling in the myocardium. **(A)** Supervised Ingenuity Pathway Analysis (IPA) of genes differentially expressed between TG^AC8^ and WT heart (n=8 per group) highlights a potential connection between ADCY8, cAMP, PKA, and RelA/NF-kB activation. **(B)** qRT-PCR analysis of *RelA, RelB, c-Rel, NF-kB1,* and *NF-kB2* transcripts in RNA extracted from left ventricular tissue of TG^AC8^ heart vs WT heart (n=6 per group, multiple unpaired t-test). **(C)** Western blot analysis of RelA, RelB, and c-Rel proteins using the left ventricular (LV) lysate of TG^AC8^ heart vs WT heart (n=6 per group, multiple unpaired t-test). **(D)** Western blot analysis of RelA protein using the nuclear lysate of LV tissue of TG^AC8^ heart vs WT heart (n=4 per group). **(E)** Representative Western blot analysis of RelA protein using the nuclear lysate of primary cardiomyocytes of the TG^AC8^ heart vs WT heart (n=4). **(F)** Representative Western blot analysis of RelA protein using the nuclear lysate of HL-1 cells transfected with GFP-AC8 or vector control (n=4 per group). **(G)** Transcription Factor activation assay for RelA protein using a nuclear lysate of HL-1 cells transfected with either GFP-AC8, shAC8, IkBm, or shRelA plasmids (n=4 per group, one-way ANOVA). **(H)** Representative Western blot analysis of IkB, pIkB, IkBKa/b, and pIkBKa/b proteins using the cell lysate of HL-1 cells transfected with GFP-AC8 (n=4 per group). **(I)** RelA ELISA-based transcription factor activation assay performed using nuclear extract of LV lysate of TG^AC8^ heart vs. WT heart (n=4, multiple unpaired t-test). **(J)** qRTPCR analysis of RelA target genes (SPP1, Gal3, CX3CL1, CXCL13, and NCAM1) in RNA extracted from LV tissue of TG^AC8^ heart vs. WT heart (n=6, multiple unpaired t-test). Graph bars represent the mean ± Standard Error of the Mean.

To confirm this, we performed additional analyses using HL-1 cells (immortalized mouse cardiomyocytes). First, we transfected HL-1 cells with a GFP-AC8 plasmid and analyzed RelA nuclear translocation via western blot. AC8 overexpression resulted in RelA nuclear translocation (Figure 6F). Next, we measured RelA concentration in HL-1 nuclear extracts using an ELISA-based Transcription Factor activation assay for RelA, and compared cells transfected with GFP-AC8 (AC8 overexpression), shAC8 (AC8 downregulation using shRNA), IkBm (expression of IkBm) or shRelA (RelA downregulation using shRNA) plasmids. Only transfection with AC8 resulted in an increase in RelA nuclear concentration (Fig. 6G). Finally, to understand whether RelA migration to the nucleus is associated with the activation of NF-κB signaling (phosphorylation of IkB and IkBKa/b proteins), we measured IkB, pIkB, IkBKa/b, and pIkBKa/b expression in nuclear lysates from the HL-1 cells transfected with GFP-AC8 or empty vector. AC8 overexpression triggered an increase in pIkB and pIKKa/b proteins expression as assessed by Immunoblot, suggesting activation of RelA signaling (Fig. 6H).

To corroborate the data obtained in HL-1 cells, we performed an Enzyme-Linked Immunosorbent Transcription Factor Activation Assay in LV extracts from WT and TG^AC8^ hearts, and quantified RelA targets in WT and TG^AC8^ mice hearts via western blot. Figure 6I shows that analysis of nuclear extracts of LV tissue from WT and TG^AC8^ mice highlighted activation of RelA in the TG^AC8^ heart vs WT (p=0.006). Figure 6J shows that the RelA targets *Spp1, Gal3, Cx3cl1, Cxcl13*, and *Ncam1* were markedly upregulated in TG^AC8^ mice hearts.

### AC8 does not bind directly to RelA and activates RelA via calcium-mediated signaling

To gain insight into the mechanistic basis of AC8-mediated RelA activation in cardiomyocytes, we first performed mass spectrometry analysis, using AC8-flag overexpression in HL-1 cells to rule out a direct interaction between RelA and AC8. Pulldown experiments using Flag antibody revealed approximately forty-six binding partners of AC8; RelA was not among these binding partners (Supplementary Figure 12, Supplementary Table 5). Since IPA analysis (Figure 6A) suggested a potential connection between cAMP signaling/PKA signaling and RelA, and our previous work revealed higher levels of *Serca2a, Ncx-1, Ltcc, CamkII,* and *Irak4* in the LV tissue of TG^AC8^ mice^7^, we tested the idea that AC8 might regulate RelA nuclear translocation through calcium signaling. Figure 7 shows that nuclear migration of RelA into the nucleus in cardiomyocytes from TG^AC8^ mice was significantly reduced by treatment for 6 hours with the cell-permeant specific calcium chelator BAPTA-AM, the Ryanodine receptor inhibitor Ryanodine, which locks the ryanodine receptor in a sub-conductance state and thus depletes the sarcoplasmic reticulum calcium store, and the PKI inhibitor peptide PKI 14-22 amide, which inhibits calcium/calmodulin-dependent Protein Kinase signalling^16, 17^. In contrast, exposure for 6 hours to 200 µM calcium chloride increased the migration of RelA into the nucleus of cardiomyocytes (Fig. 7). These results suggest that AC8 overexpression stimulates RelA nuclear translocation via upregulation of calcium-dependent PKA-mediated signaling in the TG^AC8^ heart.

**Figure 7.**
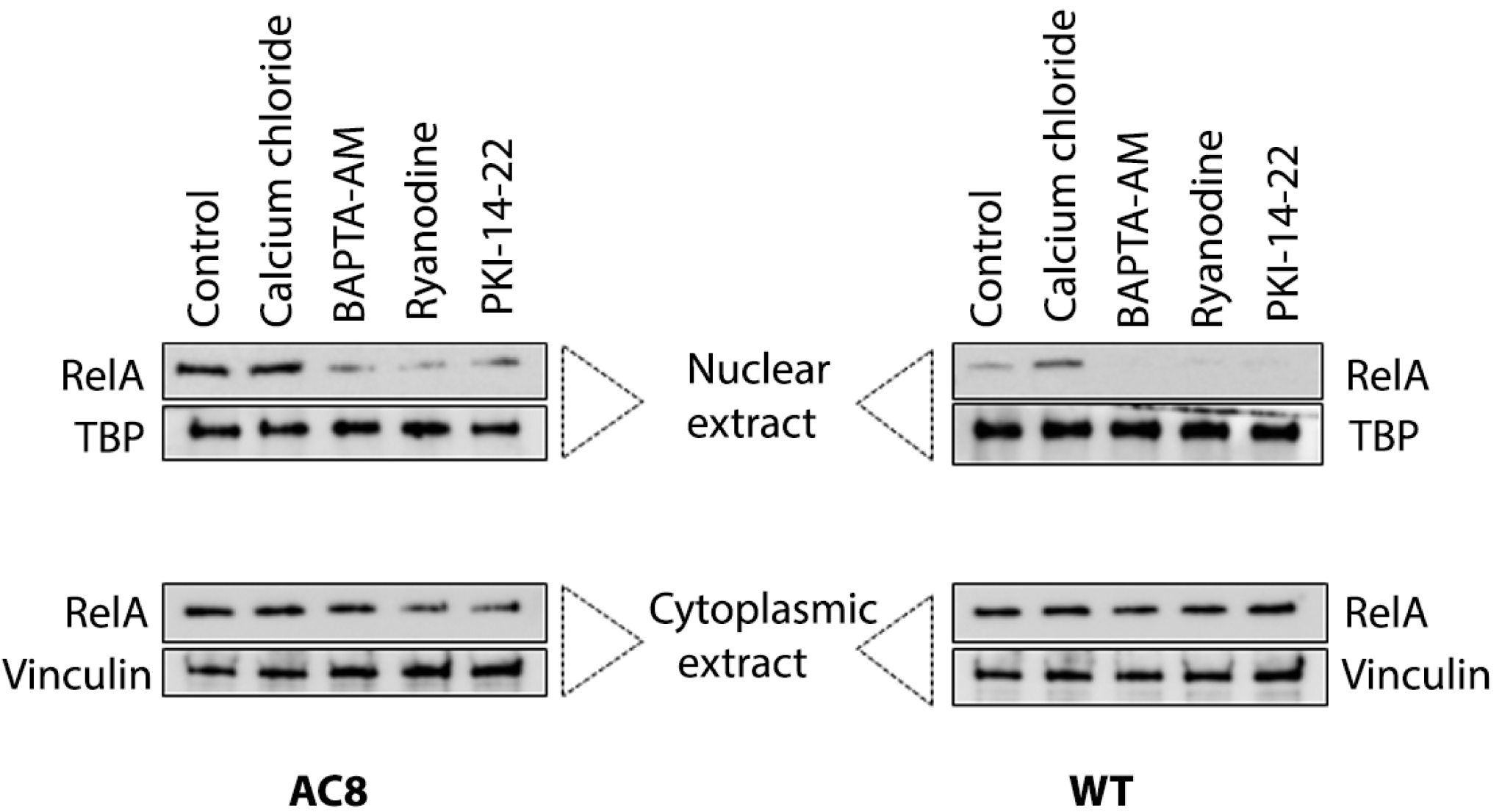
AC8 overexpression activates RelA via calcium-mediated/PKA-mediated signaling. Representative Western blot analysis of RelA protein using the cytoplasmic and nuclear lysates of cultured primary cardiomyocytes harvested from the heart of TG^AC8^ and WT mice and treated with either calcium chloride (200uM for 6 hours), BAPTA-AM (50µM for 6 hours), Ryanodine (2.8nM for 6 hours), or PKI-14-22 (PKA inhibitor peptide, 1µM for 6 hours)) vs. untreated cells. After these treatments, total protein was harvested from the primary cardiomyocytes and then blotted for specific antibodies. The experiment was repeated in triplicate.

## DISCUSSION

Our results demonstrate that cardiac-specific AC8 overexpression in young TG^AC8^ mice induces myocardial and systemic inflammation (Figures 1-4), as well as broad activation of inflammation-related and fibrosis-related pathways in multiple myocardial cell types (Figure 5, supplementary figures 3-11). Upregulation of AC8 activity in cardiomyocytes results in the activation of RelA-mediated signaling in cardiomyocytes (Figure 6), a process that is likely driven by calcium-dependent/PkA-mediated signaling (Figure 7). Since the TG^AC8^ mouse is a model of chronic myocardial stress and accelerated aging, our findings provide insights into the relationship between cardiac stress, inflammation, and aging, and point to novel potential targets for the immunomodulatory-based treatment of age-associated myocardial degeneration.

In 2008 the immunologist Ruslan Mezdithov postulated that, in response to cellular stress, tissues could elicit inflammatory responses that aim at restoring homeostasis. He defined this tissue-level inflammatory response as para-inflammation, and he and others have pointed to chronic, unresolved, para-inflammation as a major driver of tissue damage in aging organisms (this has also been referred to by other authors as “inflammaging”)^18, 19, 20^. We believe that Mezdithov’s theory of para-inflammation explains our findings and highlights their significance. In the TG^AC8^ mouse, in fact, cardiomyocytes are under constant stimulation from a constitutively active adenylyl cyclase 8. We have recently detailed how this stimulation results in a constant state of cellular stress that, in young (3 months old) mice, is associated with the activation of a remarkable array of adaptive responses^7^. Here we detailed the fact that this adaptive response includes marked activation of an inflammatory process, and it is, therefore, a “para-inflammatory” response.

There is little data on how myocardial immune cells change with aging. However, the available literature indicates that myocardial aging is associated with an increase in myocardial T cells, myocardial B cells, and myocardial granulocytes, and a decrease in myocardial monocytes/ macrophages^19^. In the young TG^AC8^ mouse, we observed an increase in all immune cell lineages, including monocytes and macrophages. The “pattern” of immune infiltration observed in the AC8 mouse, therefore, is in line with what is observed in normal aging, but has unique features. We think that also this difference can be explained using Medzhitov’s theory of para-inflammation^20^. Medzithov, in fact, postulated that para-inflammation could take different forms depending on the degree of the problem that is experienced^20^. We speculate that the overexpression of AC8 in cardiomyocytes replicates the state of cellular stress that cardiomyocytes experience during physiological aging but pushes it to a higher degree of cellular stress than that observed in physiologic aging. Therefore, the TG^AC8^ myocardium reaches a higher degree of para-inflammation than the normal aging myocardium, and experiences infiltration of monocytes and macrophages in addition to the infiltration of lymphocytes and neutrophils typically observed with aging^19^.

In the TG^AC8^ heart, together with changes in the composition of the myocardial leukocyte pool, we observed the activation of myocardial cells other than cardiomyocytes. The most remarkable findings arguably came from the analysis of endothelial cells, which showed upregulation of endothelin signaling (a marker of endothelial senescence^21^) and activation of signaling pathways related to fibrosis and cardiac hypertrophy, that are some of the key markers of age-associated cardiac degeneration^22^. Cardiomyocytes specific upregulation of AC8 was also associated with systemic inflammatory changes, as defined by changes in serum levels of inflammatory cytokines and changes in spleen size and in the immune composition of the bone marrow, and lymph nodes. In the context of the systemic activation of the immune system observed, we speculate that the reduction in spleen size observed in AC8 animals might be the result of aggressive mobilization of splenic immune cells.

In physiological aging, it is impossible to separate the aging of the heart from the aging of other organs and the aging of the immune system, and therefore it is impossible to investigate the direct effect of myocardial para-inflammation on the immune system in extracardiac compartments. Our analysis of the young TG^AC8^ mouse suggests that para-inflammation in the myocardium is sufficient to elicit systemic inflammatory changes. This is very interesting but not completely unexpected. In fact, it is well appreciated that myocardial immune cells recirculate between the myocardium, the spleen, and other organs, both at baseline and in the context of myocardial injury^23–26^.

It bears emphasis that in the TG^AC8^ hearts we found inflammation but not fibrosis. Old TG^AC8^ mice have been shown to have increased myocardial fibrosis ^6^. Therefore, we hypothesize that chronic presence of inflammation, in the context of aging, eventually results in myocardial fibrosis.Seeking to identify the mechanistic basis of myocardial para-inflammation, we performed a supervised analysis of our transcriptomic data which signaled that cAMP-mediated activation of NF-κB might play a pivotal role in this process. As NF-κB expression is upregulated in the aged rat heart^,28^, pursuing the clues provided by the “omics” analysis, we found that AC8 overexpression is associated with the nuclear translocation of RelA, a master regulator of the expression of a multitude of pro-inflammatory genes^29^. It stands to reason that chronic activation of RelA-mediated signaling would result in the secretion of inflammatory cytokines, activation of endothelial cells and smooth muscle cells, recruitment of multilineage immune cells, and eventually activation of a systemic inflammatory response. All of this is likely part of the circuitry that is activated in the 3-month-old TG^AC8^ mouse as a protective mechanism, but has the potential to become maladaptive with prolonged, persistent activation.

Nuclear translocation of RelA and activation of RelA-mediated signaling can be induced by many different stimuli, including cAMP-mediated signaling and PKA-dependent signaling^9, 29, 31,32^. We have recently shown that the 3-month-old TG^AC8^ young heart is characterized by marked activation of the adenylate cyclase/PKA/Calcium signaling^7,33^. We, therefore, assessed the requirement of PKA-dependent calcium signaling for the nuclear translocation of RelA. We observed indeed that in freshly isolated left ventricular cardiomyocytes from TG^AC8^ mice RelA nuclear translocation is a calcium/PKA-dependent event. This is supported by the observation that nuclear translocation of RelA was promoted by calcium supplementation and was inhibited by treatment with the Ca-Chelator BAPTA-AM, as well as by treatment with Ryanodine (which blocks the Ryanodine receptors in an open state and thus depletes the sarcoplasmic reticulum of calcium) and by the specific PKA inhibitor PKI-14-22.

Our work has several limitations that should be kept in mind when interpreting our findings. First of all, we did not generate myocardial specific RelA knockout AC8 animals. Therefore, the extent to which RelA mediated AC8-induced inflammation remains to be verified. Second, our mechanistic analysis of the molecular mechanisms connecting Adenylyl cyclase activity with PKA and RelA activation relied on chemical inhibitors and not on genetic manipulations. Third, we did not inhibit PKA in vivo, and therefore the role that PKA plays in vivo as a downstream mediator of adenylyl-cyclase-mediated inflammatory signaling remains unclear.

## CONCLUSIONS

We have previously shown that in the young TG^AC8^ heart, activation of Ca-PKA-mediated signaling pathways is associated with activation of a remarkable set of adaptive responses that allow cardiomyocytes to cope with the stress induced by chronic adenylyl cyclase activation^7^. Here we show that these same signaling pathways result in widespread myocardial and systemic activation of the immune system. These findings from a model of “accelerated aging” support the notion that activation of inflammatory pathways in the aging heart is part of an adaptive biological response induced by age-associated cellular stress ^29, 32^. In addition, this study suggests that chronic RelA-driven para-inflammation in cardiomyocytes might be a determinant of myocardial and possibly extra-myocardial senescence. Further studies will be needed to confirm this model and explore the potential role of RelA-modulating therapies to prevent and treat age-associated cardiac degeneration.

## ACKNOWLEDGMENTS

The authors would like to acknowledge Sylvie Rousseau for her assistance with completing specific experiments. This study was supported by the Intramural Research Program of the NIH, National Institute of Aging (USA), and by NHLBI grants 5K08HLO145108-03 and 1R01HL160716-01 to L.A.

## AVAILABILITY OF DATA AND MATERIALS

Any data not already reported in the manuscript or in the supplementary material is available from the corresponding authors upon reasonable request.

## METHODS

### Animal models

Mice were bred and maintained at the National Institutes of Health, National Institutes on Aging. All experimental procedures were done following the Guide for the Care and Use of Laboratory Animals published by the National Institutes of Health (NIH Publication no. 85-23, revised 1996). The experimental protocols were approved by the Animal Care and Use Committee of the National Institutes of Health (protocol # 466-LCS-2022). The TG^AC8^ mice, generated by ligating the murine α-myosin heavy chain promoter to a cDNA coding for the human AC8 gene, were a gift from Nicole Defer/Jacques Hanoune, Unite de Recherches, INSERM U-99, Hôpital Henri Mondor, F-94010 Créteil, France. All the mice were of C57/B6 background. Male three-month-old TG^AC8^ and wild-type (WT) littermates mice used for this study were provided with ad libitum access to water and a chow diet. All studies were performed with the approval of the IACUC of the National Institute of Aging.

### RNA sequencing and analysis

Hearts were harvested from 3 months old WT and TG^AC8^ mice. Total RNA from left ventricular tissue was extracted from 8 TG^AC8^ and 8 WT mice. The quality of RNAseq data was assessed using a standard NGS-QC toolkit as described in our earlier study^7^. Briefly, 75 bp single-end reads generated 30 to 40 million reads per library. Raw RNA sequencing (RNAseq) reads were aligned, trimmed, and mapped to the UCSC mm10 mouse reference genome and cDNA of the human AC8 gene and assembled using Tophat v2.0 to generate BAM files for each sample. Cufflinks v.2.1.1 was used to calculate FPKM (Fragments per Kilobase of transcript per Million mapped reads) for each sample. Differential gene expression analysis was performed with a Cuffdiff package (Cufflinks v2.1.1). Cutoff values of fold change >1.35 and FDR<0.05 were used to select differentially expressed genes between TG^AC8^ and WT groups.

### Proteome

Protein was extracted from the left ventricular tissue of the heart of 4 TG^AC8^ and 4 WT mice. The sample processing and fractionation and LC-MS/MS analysis are described in our earlier study^7^. Briefly, after determining the protein concentration, sample quality was assessed using NuPAGE® protein gels stained with fluorescent SyproRuby protein stain (Thermo Fisher). The detergents and lipids were removed using a standard methanol/chloroform extraction protocol (sample: methanol: chloroform: water– 1:4:1:3). Proteins were digested using trypsin/ Lys-C mixture (Promega) and desalted on a 10 x 4.0 mm C18 cartridge (Restek, cat# 917450210) using Agilent 1260 Bio-inert HPLC system with a fraction collector. The samples and one reference sample were labeled with 10-plex tandem mass spectrometry tags (TMT) using the standard TMT labeling protocol (Thermo Fisher). To control labeling efficiency and overall instrument performance, 200 femtomoles of bacterial beta-galactosidase digest (SCIEX) were spiked into each sample before TMT labeling. Labeled peptides generated from 10 different TMT channels were combined and fractionated. High-pH RPLC fractionation was performed using Agilent 1260 bio-inert HPLC system.

The dried and desalted peptide fractions were stored at −80°C until LC-MS/MS analysis. The raw data files acquired were converted to Mascot generic format using MSConvert (http://proteowizard.sourceforge.net) as described in our earlier study^7^. The median fold change of all unique peptides was used for calculating the fold change between genotypes for each expressed protein. Genotype differences were compared using Student’s t-test.

### Complete blood count

A blood sample from 3 months old TG^AC8^ and WT mice was collected in heparin-coated tubes and shipped to the company Comparative Clinical Pathology Services, LLC for analysis. The data were analyzed using software GraphPad Prism software (Prism, version 7.0).

### Quantibody inflammation array

Plasma samples from 3 months old TG^AC8^ and WT mice were collected after the sacrifice of mice and stored at −80^0^C until analysis. The assay was performed using the Quantibody® Mouse Inflammation Array kit (QAM-INF-1-1, RayBiotech Inc.). The samples were diluted 4x using the sample diluent. 100 ul of both samples and standards were loaded onto the glass slide (labeled with the cytokines and chemokines) and incubated for 2 hours. Samples and standards were decanted from each well and washed with 150 µl of 1X Wash Buffer I at room temperature. The detection antibody cocktail was added to each well and incubated at room temperature for 1 hour. The samples were decanted and washed with 150 µl of 1X Wash Buffer I at room temperature. A Cy3 equivalent dye-conjugated streptavidin was added to each well and incubated in the dark at room temperature for 1 hour. The samples were decanted from each well, washed with 1x wash buffer, and dried. The slide was imaged for fluorescent signals using a laser scanner equipped with a Cy3 wavelength (green channel). The data was extracted, computed in the standard format provided by the company, and analyzed for relative levels of different cytokines.

### Histology

Heart tissues harvested from the TG^AC8^ and WT mice were washed with ice-cold PBS and fixed overnight with 4% paraformaldehyde. The fixed heart tissues were then paraffin-embedded, and 5 µm thick sections were cut from these paraffin blocks. The heart sections were stained with Hematoxylin and Eosin or Trichrome stain and analyzed for myocardial structure and fibrosis. The captured images from heart sections were quantified using the software KEYENCE BZ-X800 analyzer.

### Immunohistochemistry

Heart tissue sections were processed for immunostaining using the BioGenex kit as per the manufacturer’s protocol. Briefly, the tissue sections were deparaffinized and treated with an antigen retrieval solution. The sections were then treated with peroxide block to mask endogenous peroxide and incubated with primary antibody overnight at 4°C. After washing twice with wash buffer, sections were incubated with Polymer-HRP for 40 minutes at room temperature. The nucleus was stained with hematoxylin; sections were mounted with coverslips and visualized under the microscope. Immunostaining was quantified using the software KEYENCE BZ-X800 analyzer.

### Troponin I assay

Plasma samples from 3 months old mice were used to measure cardiac Troponin I. Troponin I was measured with the assistance of the Johns Hopkins Hospital Clinical Laboratory using the high-sensitivity Abbot Troponin assay. Serum was diluted 1:4 prior to analysis using a diluent optimized for the assay. The assay was validated using serial dilutions of serum collected from mice 24 hours after coronary ligation and including negative controls.

### Flow cytometry

The heart was perfused with ice-cold PBS using a 25G needle, harvested, and minced. The single cell suspension was prepared by digesting the heart in HBSS supplemented with Collagenase 1 (450 U/ml), Hyaluronidase (60 U/ml), and DNase I (60 U/ml) for 30 minutes at 300rpm, 37°C. The enzymes in the cell suspension were deactivated by adding HBSS supplemented with 5% FBS. Red blood cell lysis was performed using ACK lysis buffer for 5 minutes at room temperature. Cells were then washed with HBSS containing 5% FBS and resuspended with 300 µl FACS buffer (DPBS, 2% FBS, 2 mM EDTA). Cells were stained with an antibody cocktail for 30 minutes at 4°C in the dark. The sample was washed twice and resuspended in 500 µl FACS buffer. Helix green or Live-dead aqua dye was used to exclude dead cells from the analysis. Immune cells were gated as CD45+. Monocytes were gated as CD45+ CD11b+ Ly6G-CD64-Ly6C^high^. Neutrophils were gated as CD45+CD11b+Ly6G+. B cells and T cells were gated as CD45+CD11b-CD19+, and CD45+CD11b-CD4+/CD8+, respectively. Macrophages were gated as CD11b+Ly6G-CD64+Ly6C^low^. The flow cytometric analysis was performed using BD Symphony and BD Fusion machines. The flow data were analyzed using the software FlowJo. The antibodies used for flow cytometry experiments are listed in the supplementary table 6.

### qRT-PCR

Snap-frozen left ventricular tissue was homogenized, and total RNA was isolated using RNeasy Mini Kit (Qiagen). Approximately 2µg of RNA was reverse transcribed into cDNA using reverse transcriptase (NEB), and quantitative real-time PCR was performed using SYBR green (Applied Biosystems). The specificity of the primer was determined by melting curve analysis (single product amplification). mRNA expression of the target and the control groups was normalized with GAPDH, and fold change between target and control groups was determined using the method 2^ΔΔCt^ as described previously^35^. The primers used in this study are listed in supplementary table 6.

### Western blot analysis

Snap-frozen left ventricular tissue collected after the sacrifice of mice was homogenized in RIPA buffer with protease inhibitor cocktail (Roche) using prechilled Precellys bead homogenizer. The protein concentration in each sample was determined using Bicinchoninic acid (BCA) assay. 20-30ug of protein was loaded on precast Tris-Glycine SDS-PAGE gel (BioRad) and transferred to the PVDF membrane by semidry transfer. Primary antibodies were incubated overnight in 3% BSA in TBST buffer. Enhanced chemiluminescence (ECL) signal was detected using the Amersham Imager 600 imaging system. Antibodies for immunoblot are listed in the supplementary table 6.

### TransAM p65 NF-κB Assay

The nuclear lysate was prepared from HL-1 cells or fresh LV tissue as per standard protocol (Thermo Scientific kit). TransAM p65 NF-κB activation assay was performed using nuclear lysate as per standard protocol (Active Motif). Briefly, recombinant NF-κB protein (0ng/µl −10ng/µl) as standard, positive control (TNFα stimulated Jurkat nuclear extract), and samples (nuclear extract containing activated transcription factor) were added to the oligonucleotide coated plate and incubated for 1 hour at room temperature. The plate was washed and incubated with p65 antibody for 1 hour. An HRP-conjugated secondary antibody was added to the plate and incubated for 1 hour. The colorimetric antibody binding signal was then detected using a spectrophotometer at 450nm.

### Single-cell RNA sequencing

A single cell suspension from the heart was prepared as described earlier in flow cytometry. During digestion, cells were stained with Vybeant DyeCycle Violet Ready Flow Reagent (Invitrogen), adding two drops per 3ml digestion reaction. This allowed visualization of live cells. Counterstain of dead cells was performed with propidium iodide. Cells were then stained with anti-CD45 and CD31 antibodies [1] [2] [3] and were sorted in a Moflo sorter. An equal number of FACS-sorted CD45+CD31-, CD45-CD31-, and CD45-CD31+ cells were mixed and processed to generate single-cell cDNA libraries using the 10X Genomics platform per the manufacturer’s standard protocols. Cardiomyocytes were excluded because their size makes them incompatible with high-throughput single cell sequencing. Cells were sequenced at a target read depth of 75,000 reads/cell using a HiSeq 4000 sequencer (Illumina). Single-cell counts were generated from unaligned reads by sequence alignment and de-multiplexing using Cell Ranger from 10x Genomics. Post-alignment QA/QC performed using single-cell counts suggested that none of the 10,667 cells expressed mitochondrial genes at greater than 5 percent of their total gene expression; cells were not excluded from further analysis. Normalized cell counts were generated using filter exclude features where value <= 0.0 in at least 99.9 % of the cells, CPM (counts per million), the scale factor of 10000, and log transformation. Of the 22,441 genes, 13,331 passed these criteria, and the remaining 9,110 were filtered out. Graph-based and K-based clusters were generated using 20 principal components derived from the differentially expressed genes. The cell clusters were defined using the top 10 markers enriched in each K-based cluster. Genes differentially expressed between specific clusters of interest were identified using ANOVA.

### Calcium signaling and RelA expression

Cardiomyocytes were isolated from both TGAC8 and WT mice and divided into different treatment groups, cultured for 1 hour, and washed with PBS to get rid of cell debris. The cells were then treated with either BAPTA-AM (Millipore Sigma) at 50uM for 6 hours or PKI 14-22 amide, a membrane permeable PKA inhibitor peptide (Tocris Bioscience) at 1uM for 6 hours, or Ryanodine (Tocris Bioscience) at 2.8nM for 6 hours or Calcium chloride at 500uM for 6 hours or Vehicle control (0.1% DMSO in phosphate buffer saline) for 6 hours and harvested for preparation of cytoplasmic and nuclear lysates. The nuclear expression of RelA was determined using Western blot analysis as described earlier.

### Statistical Analysis

Data were analyzed using GraphPad Prism software (Prism, version 7.0). The sample means were compared using either an unpaired t-test, multiple unpaired t-test, or a one-way analysis of variance. Multiple comparisons were performed using post hoc Tukey’s test wherever applicable. *P*<0.05 was considered a statistically significant difference. Figure legends indicate the specific statistical tests used in each analysis

## SUPPLEMENTARY FIGURES AND SUPPLEMENTARY TABLES

**Supplementary figure 1. AC8 overexpression increases myocardial inflammation.** Relative protein enrichment (LC-MS) of inflammation-related proteins (Fold change>1.5, p-value<0.05) in the LV tissue of TG^AC8^ vs WT mice (n=8 per group).

**Supplementary figure 2. AC8 overexpression induces the recruitment of multilineage immune cells to the heart. (A)** Representative gating strategy used to analyze myocardial immune cells via flow cytometry. **(B)** Representative FACS plot of CD45+ cells, CD45+ CD11b+ cells, CD19+ cells, CD19+ CD11b+ cells, CD19+IgM+IgD+ cells, CD19+IgM-IgD-cells, CD19+IgM+IgD-cells, CD19+IgM-IgD+ cells, CD4+ and CD8+ T cells of TG^AC8^ heart vs WT heart. **(C)** Representative FACS plot from the blood of TG^AC8^ mice vs WT mice.

**Supplementary figure 3. Ingenuity pathway analysis (IPA) of genes differentially expressed in smooth muscle cells from TG^AC^**^8^ **and WT hearts.** Genes with a 1.5 fold-change and p-value < 0.05 were used for pathway analysis. The graph shows the pathways with a z-score > 2 and a p-value < 0.05 in the IPA analysis.

**Supplementary figure 4. Ingenuity pathway analysis (IPA) of genes differentially expressed between fibroblasts from TG^AC^**^8^ **and WT hearts**. Genes with a 1.5 fold-change and p-value < 0.05 were used for pathway analysis. The graph shows the pathways with a z-score > 2 and a p-value < 0.05 in the IPA analysis.

**Supplementary Figure 5. Ingenuity pathway analysis (IPA) of genes differentially expressed in B cells from TG^AC^**^8^ **and WT hearts**. Genes with a 1.5 fold-change and p-value < 0.05 were used for pathway analysis. The graph shows the pathways with a z-score > 2 and a p-value < 0.05 in the IPA analysis. The results support the hypothesis of activation of the B cells by the cytokines in the microenvironment.

**Supplementary Figure 6. Ingenuity pathway analysis (IPA) of genes differentially expressed in CD8+ T cells from TG^AC^**^8^ **and WT hearts**. Genes with a 1.5 fold-change and p-value < 0.05 were used for pathway analysis. The graph shows the pathways with a z-score > 2 and a p-value < 0.05 in the IPA analysis. The results support the hypothesis of activation of the CD8+ T cells by the cytokines in the microenvironment.

**Supplementary Figure 7. Ingenuity pathway analysis (IPA) of genes differentially expressed in CD4+ T cells from TG^AC^**^8^ **and WT hearts**. Genes with a 1.5 fold-change and p-value < 0.05 were used for pathway analysis. The graph shows the pathways with a z-score > 2 and a p-value < 0.05 in the IPA analysis.

**Supplementary Figure 8. Ingenuity pathway analysis (IPA) of genes differentially expressed in Glial-like cells from TG^AC^**^8^ **and WT hearts**. Genes with a 1.5 fold-change and p-value < 0.05 were used for pathway analysis. The graph shows the pathways with a z-score > 2 and a p-value < 0.05 in the IPA analysis.

**Supplementary Figure 9. Ingenuity pathway analysis (IPA) of genes differentially expressed in macrophages from TG^AC^**^8^ **and WT hearts.** Genes with a 1.5 fold-change and p-value < 0.05 were used for pathway analysis. The graph shows the pathways with a z-score > 2 and a p-value < 0.05 in the IPA analysis. The results suggest that the macrophages could be actively extravasating to the tissue and is activated by the cytokines in the microenvironment and local immune cells.

**Supplementary Figure 10. Ingenuity pathway analysis (IPA) of genes differentially expressed in neutrophils from TG^AC^**^8^ **and WT hearts**. Genes with a 1.5 fold-change and p-value < 0.05 were used for pathway analysis. The graph shows the pathways with a z-score > 2 and a p-value < 0.05 in the IPA analysis.

**Supplementary Figure 11. Ingenuity pathway analysis (IPA) of genes differentially expressed in monocytes from TG^AC^**^8^ **and WT hearts**. Genes with a 1.5 fold-change and p-value < 0.05 were used for pathway analysis. The graph shows the pathways with a z-score > 2 and a p-value < 0.05 in the IPA analysis.

**Supplementary Figure 12. Representative Western blot analysis of Flag protein using cell lysate of HL-1 cells transfected with Flag-AC8 or vector control (n=3)**. To identify the binding partners of the AC8 protein, HL-1 cells were transfected with Flag-AC8 or vector control, and transfected cells were then immunoprecipitated using a flag antibody. Three independent Flag-ADCY8 overexpressed HL-1 cell lysates were pooled together and pulldown with antiflag/IgG antibody, run on SDS-PAGE, destained, resolved bands were cut from the SDS-PAGE gel at 55KDa, <55KDA and >55KDa and sent for MALDI-TOF_LCMS analysis. Proteins with peptide count >3 and not identified in IgG or unstained SDS-PAGE gel samples were included in the analysis.

**Supplementary Table 1. RNA-seq and proteome (LC-MS) analysis show differentially expressed genes and proteins between the TG^AC^**^8^ **and WT heart**. The table shows the raw gene expression and raw protein expression files.

**Supplementary Table 2. Ingenuity Pathway Analysis (IPA) shows upstream regulators of gene expression differences observed between the TG^AC^**^8^ **and WT heart**. Genes with a 1.5 fold-change and p-value < 0.05 were used for pathway analysis. The list of differentially expressed genes and the list of upstream regulators identified by IPA are reported.

**Supplementary Table 3. Quantibody inflammation arrays of plasma samples of TG^AC^**^8^ **and WT mice (n=4 per group)**. Plasma was collected from 4 TG^AC^^8^ and 4 WT mice and analyzed using quantibody inflammation array kit (Sandwich-based assay). The table shows the cytokine concentration (pg/ml) of 40 different inflammatory cytokines analyzed using this kit.

**Supplementary Table 4. Complete blood count analysis of TG^AC^**^8^ **and WT mice (n=6 per group)**. The table shows the absolute no. (k/ul) and % of white blood cells (WBCs), neutrophils (NE), Lymphocytes (LY), Monocytes (MO), Eosinophils (EO), Basophils (BA), Nucleated red blood cells (NRBCs), and red blood cells (RBCs). The table also shows the hemoglobin (HB), hematocrit (HCT), mean corpuscular volume (MCV), Mean corpuscular hemoglobin (MCH), Mean corpuscular hemoglobin concentration (MCHC), red blood cell distribution width (RDW), reticulocyte count (retics) and platelet count (PLT) of TG^AC^^8^ and WT mice.

**Supplementary Table 5**. **MALDI-TOF-LC-MS analysis shows binding partners of AC8 protein from HL-1 cells transfected with Flag-AC8 (n=3)**. To identify the binding partners of the AC8 protein, HL-1 cells were transfected with Flag-AC8 or vector control, and transfected cells were then immunoprecipitated using a flag antibody. Three independent Flag-AC8 overexpressed HL-1 cell lysates were pooled together and pulldown with antiflag/IgG antibody, run on SDS-PAGE, destained, resolved bands were cut from the SDS-PAGE gel at 55KDa, <55KDA and >55KDa and sent for MALDI-TOF_LCMS analysis. Proteins with peptide count >5 and not identified in IgG or unstained SDS-PAGE gel samples were included in the analysis (Worksheet 1 and worksheet 2).

**Supplementary Table 6. A list of primers and antibodies used in this study.**

